# The dynamics of speciation in the gall-forming aphid *Geoica* spp. within and among *Pistacia* host tree species

**DOI:** 10.1101/2022.03.20.485014

**Authors:** Rachel Ben-Shlomo, Stav Talal, Moshe Inbar

**Affiliations:** Department of Biology and Environment, Faculty of Natural Sciences, University of Haifa-Oranim, Tivon, 36006, Israel; Department of Evolutionary and Environmental Biology, Faculty of Natural Sciences, University of Haifa, Haifa, 3498838, Israel

**Keywords:** *Geoica spp.*, gall-forming aphids, haplotype, speciation

## Abstract

Trees of the genus *Pistacia* serve as obligate hosts for gall-forming aphids (Hemiptera, Aphididae, Fordini). Each aphid species induces a characteristic gall on a single *Pistacia* host species. The genus *Geoica* (Fordini) induce similar spherical closed galls on the lower side of the leaflet’s midvein, on different *Pistacia* species. Two species of *Pistacia* trees that harbor *Geoica* galls grow naturally in Israel: *P*. *palaestina* and *P. atlantica.* We analyzed the phylogeny and the genetic structure of the *Geoica* species complex in Israel, and assessed the genetic differentiation and the level of host plant specificity of the aphids between *P*. *atlantica* and *P. palaestina*. We found that the splitting of the genus between *P. atlantica* and *P. palaestina* is estimated to have occurred 24–25 Ma (the Oligocene/Miocene boundary). Five different haplotypes suggesting five different species have been further speciating among *Geoica* spp., galling on *P. atlantica*, and an additional three species, on *P. palaestina*.

**Accession Numbers (COI sequences):**

MW452548 - Pp-Q-7d-Gamla - Haplotype-VI
MW452549 - Pp-Q-10b-Gamla - Haplotype-VII
MW452550 - Pp-A-1a-Mezar - Haplotype-VII
MW452551 - Pp-S-6c-Sasa - Haplotype-VI
MW452552 - Pp-New Hermon-2-5 - Haplotype-VIII
MW452553 - Pa-Q-10a-II Gamla - Haplotype-IV
MW452554 - Pa-Q-5d Gamla - Haplotype-II
MW452555 - Pa-A-2f Mezar - Haplotype-IV
MW452556 - Pa-E-13 Anafa - Haplotype-III
MW452557 - Pa-K-10a Adolam - Haplotype-IV
MW452558 - Pa-K-3a Adolam - Haplotype-II
MW452559 - Pa-N-2a Nizana - Haplotype-I
MW452560 - Pa-Z-1-3 Hazeva - Haplotype-I
MW452561 - Pa-SL-3-3 Jordan-Sela - Haplotype-V

## 1. INTRODUCTION

Galls are deformed growths of plant tissues that are induced by numerous organisms, primarily insects. The formation of galls requires close associations between the inducers and the host plants. Indeed, most gall-forming insects are strictly specific to a given host plant species, organ and tissue. The mechanism of gall formation and maintenance, and the way insects modify plant developmental pathways is still unknown (Giron et al., 2016). However, typically, the morphology of the galls is associated with the inducing insect species.

Trees of the genus *Pistacia* serve as obligate hosts for gall-forming aphids (Hemiptera, Aphididae, Fordini) (Wool, 2004). The galls are formed early in the spring by a single aphid (the fundatrix), which feeds on the phloem sap. Within each gall several additional generations are produced parthenogenetically (females only). In the fall, winged aphids (the fall alatae) leave the galls and migrate to the roots of non-specific herbaceous host plants (Wool, 2004).

Each aphid species induces a characteristic gall on a single *Pistacia* host species. The galls have distinct shapes, colors and sizes (Inbar et al., 2004; Koach and Wool, 1977; Wool and Burstein, 1991). In most cases, the shape of the gall is characteristic of a particular species of aphid, and as such, can also be used as a “tool” in *Pistacia* systematics (Inbar, 2008), as in other hosts of gall-forming insects (Aguilar and Boecklen, 1992; Floate et al., 1996).

The genus *Geoica* (Fordini) represents an exceptional uncoupling between gall morphology and aphid-species identity. *Geoica* is a Palaearctic genus that forms galls on *Pistacia* trees and alters the host in the fall, to roots of Poaceae, and sometimes Cyperaceae (Blackman and Eastop, 2018). Most *Geoica* species (about 10 species) induce similar spherical (ball-shaped) closed galls, located on the lower (abaxial) side of the leaflet’s midvein, on different *Pistacia* species.

The taxonomic status of the genus *Geoica* is challenging and unclear. Based on the morphometry of fall alatae, Brown and Blackman (1994) analyzed the so-called *G. urticularia* complex. They divided the complex into five species inhabiting the Mediterranean parts of Europe, North Africa, the Levant and Iran: *G. rungsi* and *G. harpazi* on *P. atlantica; G. muticae* on *P. mutica; G. urticularia* on *P. terebinthus*; and *G. wertheimae* on *P. palaestina*. These species were not associated with any given gall trait. Limited molecular analyses suggested there were three species of the genus *Geoica* in Israel, *G. wertheimae* on *P. palaestina*, and two putative species on *P. atlantica*, both collected in northern Israel (Inbar et al., 2004). These two species, which may possibly be *G. rungsi* and/or *G. harpazi* (Brown and Blackman, 1994), were marked by Inbar et al. (2004) as *Geoica* sp. An additional species producing spherical galls, found on the leaf petiole (not on the leaflet midvein) of *P. atlantica* trees, which was identified at the time as *G. swirskii* (Remaudiere et al., 2004), has been assigned to a new genus, *Inbaria* (Barjadze et al., 2018). Blackman and Eastop (2018) recently revised the status of the *G. utricularia* complex. Considering the substantial variation in the morphology of the galls and of the emigrant alatae and their embryos, they suggested additional species, as yet unnamed, in the genus *Geoica*; these species are closely related to *G. utricularia*, but associated with other *Pistacia* spp.

Two species of *Pistacia* trees that harbor *Geoica* galls grow naturally in Israel: *P. palaestina* and *P. atlantica*. The former is common in Mediterranean habitats, while the latter is found in the Mediterranean and Irano-Turanian distribution zone. Both species inhabit diverse habitats along climatic gradients that vary in solar radiation, temperature and precipitation. The origin and evolution of the genus *Pistacia* are controversial (Kozhoridze et al., 2015; and references therein); hence, most studies suggested that the genus evolved more than 80 M years ago. Within the genus *Pistacia, P. palaestina* and *P. atlantica* are placed in different phylogenetic lineages (e.g., Golan-Goldhirsh, 2009; Kafkas and Perl-Treves, 2002; Zohary, 1972; Xie et al., 2014). Nevertheless, natural crossing between the two species does exist (Golan-Goldhirsh, 2009). Natural populations of *P. palaestina* and *P. atlantica* are found sympatrically in several regions in northern and central Israel, while other populations are clearly allopatric. Galls of the genus *Geoica* can be readily found throughout the distribution zone of both *Pistacia* tree species in Israel, including on hybrid trees of *P. atlantica x P. palaestina* (Ben-Shlomo and Inbar, personal observations).

The East Mediterranean region, at the margin of the Eurasian and African plates, went through massive geological changes during the late Tertiary (66 to 2.6 Ma) and the succeeding Quaternary (Pleistocene) glaciation. These changes involved multiple events of land contacts and withdrawals (Lymberakis et al., 2007; and references therein). Such infrequent connections possibly facilitated occasional openings for species dispersal between the Irano-Turanian and the Mediterranean distribution zones.

The complex systematics and taxonomy of the genus *Geoica*, and its unique and fragmented distribution on its *Pistacia* host species (and hybrids), suggest that it may have speciated in the Middle East, in general, and in Israel, in particular, where we find allopatric and sympatric populations across different climatic regions. The main goal of this study was to trace the speciation processes of the genus *Geoica* in the Levant by analyzing the phylogeny and genetic structure of the species complex in Israel, using several molecular markers (nuclear microsatellite loci and sequences of the mitochondrial COI and 12S genes). Specifically we assessed: 1. the genetic differentiation of the aphids (both between and within populations) between *P. atlantica* and *P. palaestina*, as well as the level of host plant specificity (i.e., are particular *Geoica* aphids in sympatric populations found solely on a specific host *Pistacia* species?); 2. the genetic diversity of the *Geoica* aphids between populations within each species; 3. the association between gall morphology and aphid species on both *Pistacia* hosts; and 4. the timeline of speciation according to the molecular clock, if speciation came about.

## 2. MATERIAL AND METHODS

### 2.1 Sampling

At the end of the summer (September–October) *Geoica* galls were collected throughout their distribution range of *P. atlantica* and *P. palaestina* in Israel. In addition, we had several samples collected from *P. atlantica* trees in various localities in Moab and Edom in Jordan (Fig. 1 and Table 1). The sampling (in total 141 galls) included allopatric and sympatric populations: 59 galls from *P. palaestina*, 77 from *P. atlantica* and 5 galls from a single hybrid tree (*P. atlantica* x *P. palaestina*).

**Figure 1:**
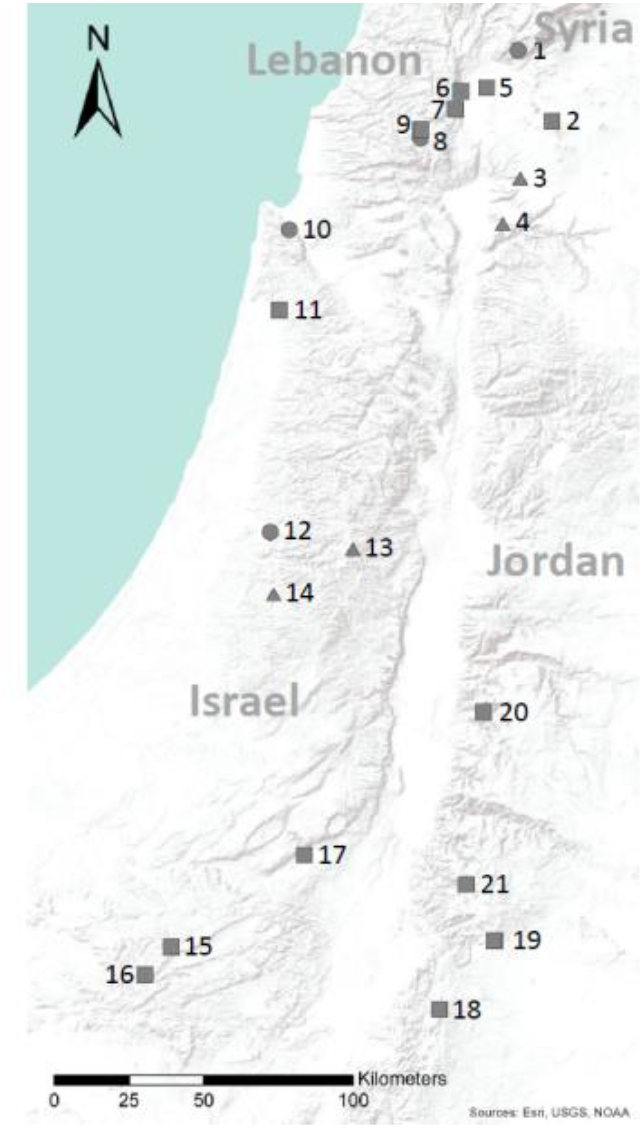
Distribution map and sampling sites in Israel and Jordan of *P. atlantica* (square), *P. palaestina* (circle) and sympatric populations of *P. atlantica* and *P. palaestina* (triangle).

**Table 1:** Sampling of *Geoica* galls throughout the range of *Pistacia atlantica* and *P. palaestina* in Israel and Jordan. Numbers are following Fig 1.

| Country | Number | Population | *Pistacia*<br>*atlantica* | *Pistacia*<br>*palaestina* | *Hybrid*<br>*palaestina x*<br>*atlantica* |
| --- | --- | --- | --- | --- | --- |
| Israel | 1 | Mt Hermon |  | 16 |  |
|  | 2 | Hazeka | 1 |  |  |
|  | 3 | Gamla | 8 | 11 |  |
|  | 4 | Meizar | 2 | 2 |  |
|  | 5 | Tel Anafa | 6 |  |  |
|  | 6 | Ein Avazim | 2 |  |  |
|  | 7 | Ramat Naftali | 3 |  |  |
|  | 8 | Sasa |  | 13 |  |
|  | 9 | Dishon |  |  | 5 |
|  | 10 | Nof Carmel |  | 2 |  |
|  | 11 | Ada | 2 |  |  |
|  | 12 | Park Canada |  | 3 |  |
|  | 13 | Jerusalem | 7 | 1 |  |
|  | 14 | Adolam | 10 | 11 |  |
|  | 15 | Nizana | 10 |  |  |
|  | 16 | Lotz | 4 |  |  |
|  | 17 | Hazeva | 11 |  |  |
| Jordan | 18 | Bir Dabarat | 1 |  |  |
|  | 19 | Ja'far al-Tayyar | 3 |  |  |
|  | 20 | Hemed | 2 |  |  |
|  | 21 | Sela | 5 |  |  |
| **Total** |  |  | 77 | 59 | 5 |

### 2.2 Gall morphology

All galls are globular, completely sealed, and located on the abaxial side of the leaflet’s midvein. The size, shape and surface configuration of the galls vary considerably both within and among their host tree populations. The morphology of the galls was categorized according to the gall’s surface and shape characteristics: A. surface: smooth, partially crumpled and coarse; B. shape: ball-like, flattened discus-like, pear-like, “bumpy” spheres and coral-like (see photos in Table 2A). Gall characteristics were then associated with host plant species and the aphid’s genetic identity.

**Table 2:**
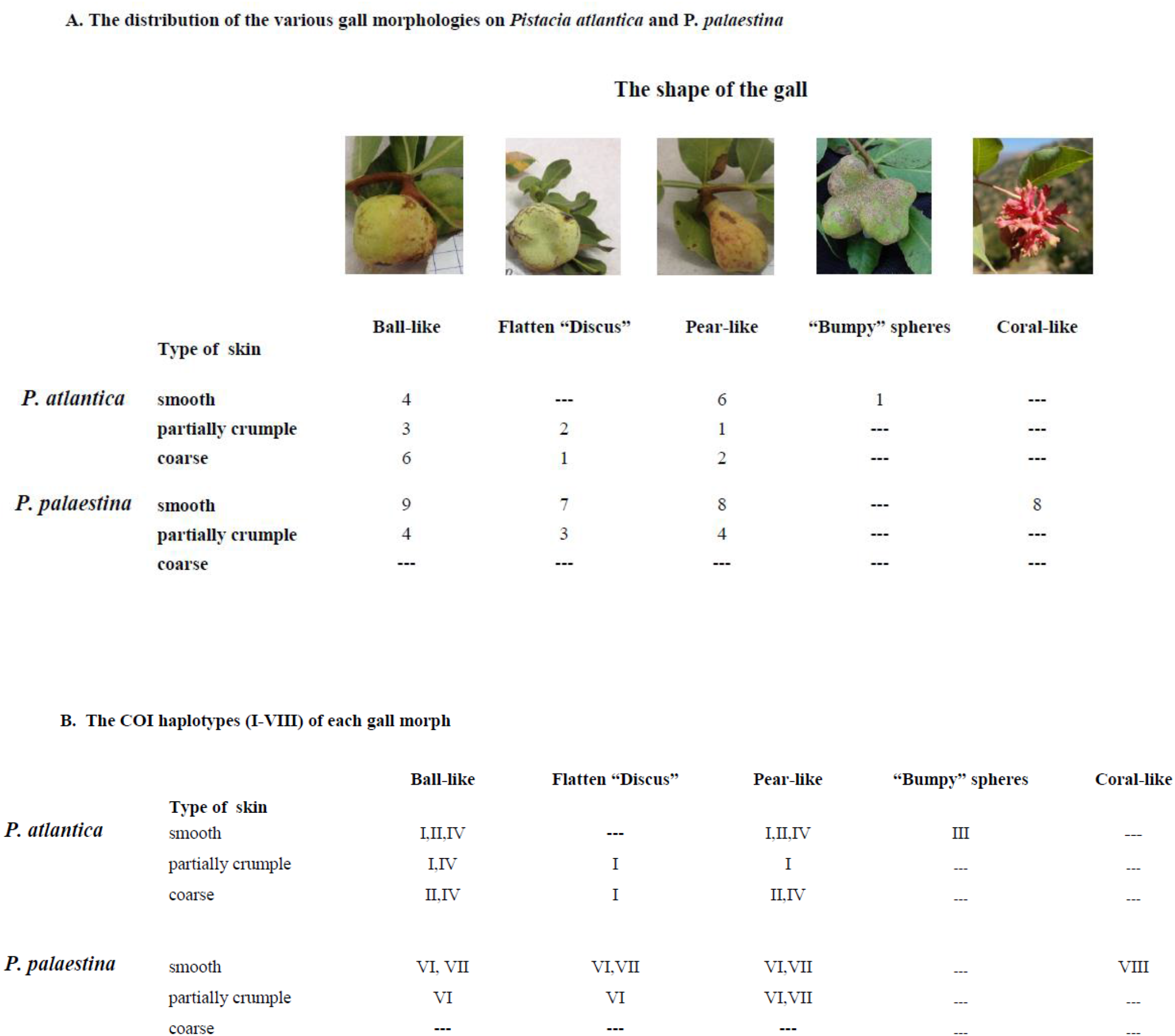
A. The distribution of the various gall morphologies on *Pistacia atlantica* and *P. palaestina*

|  |  | The shape of the gall |  |  |  |  |
| --- | --- | --- | --- | --- | --- | --- |
| Type of skin |  | Ball-like | Flatten "Discus" | Pear-like | "Bumpy" spheres | Coral-like |
| *P. atlantica* | smooth | 4 | --- | 6 | 1 | --- |
|  | partially crumple | 3 | 2 | 1 | --- | --- |
|  | coarse | 6 | 1 | 2 | --- | --- |
| *P. palaestina* | smooth | 9 | 7 | 8 | --- | 8 |
|  | partially crumple | 4 | 3 | 4 | --- | --- |
|  | coarse | --- | --- | --- | --- | --- |

B. The COI haplotypes (I-VIII) of each gall morph
|  |  | Ball-like | Flatten "Discus" | Pear-like | "Bumpy" spheres | Coral-like |
| --- | --- | --- | --- | --- | --- | --- |
| *P. atlantica* | Type of skin |  |  |  |  |  |
|  | smooth | I,II,IV | --- | I,II,IV | III | --- |
|  | partially crumple | I,IV | I | I | --- | --- |
|  | coarse | II,IV | I | II,IV | --- | --- |
| *P. palaestina* | smooth | VI, VII | VI,VII | VI,VII | --- | VIII |
|  | partially crumple | VI | VI | VI,VII | --- | --- |
|  | coarse | --- | --- | --- | --- | --- |

### 2.3 Genetic analysis of the aphids

Due to their parthenogenetic reproduction, by the fall each gall may contain up to several hundred genetically identical aphids (Wool and Ben-Zvi 1998). DNA was extracted from the pool of aphids in each gall using the DNeasy Blood and Tissue Kit (Qiagen). The sequence of fragments of the mitochondrial genes *Cytochrome Oxidase I* (COI) and the ribosomal 12S subunits, as well as analysis of seven polymorphic microsatellite loci (Ivens et al., 2010), were used as molecular markers.

#### 2.3.1 *Mitochondrial COI* and *12S sequencing*

A fragment of 630bp of the mitochondrial COI gene (n=114 galls) and a fragment of 548bp of the mitochondrial 12S gene (n=78 galls) were amplified by PCR in a total volume of 13 μ1 and at an annealing temperature of 55°C. The design of primers followed COI sequences found in the GenBank (AY227075; F-GGTCTGGGATAATTGGTTCTTC, and R-AGGCCAATT GTTATTATAGC), and arthropods-conserved 12S primers (Kocher et al., 1989; 12Sai-AAACTAGGATTAGATACCCTATTAT and 12Sbi-AAGAGCGACGGGCGATGT GT). The single PCR products of the amplified fragments were purified (Illustra ExoProstar) and sequenced using the BigDye Terminator sequencing procedure (Applied Biosystems).

The sequences’ chromatograms were visualized, checked manually, and the least clear ends of each sequence (up to 40bp from each end) were excluded from the analysis. Sequences were aligned using the software MEGA 5.1 (Tamura et al., 2011). The chromatographs of all Single Nucleotide Polymorphism (SNPs) were re-checked visually, and the resulting phylogenetic dendrograms were clustered by the Maximum Likelihood method (1,000 bootstrap).

The COI phylogeny tree was tested for molecular clocks throughout the whole data set, with the published *Baizongia pistaciae* (AY227079; Inbar et al. 2004) and *Slavum wertheimae* COI sequences (northern populations — GU391571; Negev Mountains — GU391572; Avrani et al., 2012) as outgroups. The Maximum Likelihood value for the given topology, with and without the molecular clock constraints, were compared using the Tamura-Nei (1993) model method (conducted in MEGA5.1; Tamura et al., 2011), where the null hypothesis tested was an equal evolutionary rate throughout the tree.

#### 2.3.2 Microsatellites Analysis

123 individuals from various populations and species of *Pistacia* were analyzed by seven known nuclear microsatellite loci (Gu-1, 3, 4, 6, 8, 11 and 13; Ivens et al., 2010). The F-primer of each microsatellite was labeled with fluorescent dye (6-Fam, Vic, Ned or Pet) and the amplification products were read by ABI 3130xl Fluorescence Reader (Applied Biosystems).

Microsatellites’ allele identification, genotyping and observed heterozygosity (Ho) were determined directly from the chromatographs using 500-Liz standard marker and Peak Scanner software (Applied Biosystems). Expected heterozygosity (He) and genetic identity (I) were calculated following Nei (1978). Data were analyzed by GenAlEx version 6.5b4 software (Peakall and Smouse, 2006). The analysis of molecular variance (AMOVA) procedure followed the methods of Michalakis and Excoffier (1996) using GenAlEx 6.5b4 software (999 permutations; Peakall and Smouse, 2006). Population clustering was performed using the Bayesian clustering method (STRUCTURE), which divided the samples into possible homogenous groups (subpopulations) according to their degree of genetic similarity (STRUCTURE 2.3.4 admixture model; burn-in of 100,000 steps and 100,000 iterations; five replicates for each K) (Pritchard et al., 2000, 2010). The inference of the probable number of clusters was extracted by the log likelihood for each putative number of populations (K), Ln P(D) = L(K), and by the delta K methods (Evanno et al., 2005), using the program Structure Harvester (Earl and vonHoldt, 2012).

## 3. RESULTS

### 3.1 Gall morphology

Gall morphology was typified for 43 galls collected from *P. palaestina* and 26 from *P. atlantica* (Table 2). Ball-, flattened discus- and pear-like galls were found on both *Pistacia* species, whereas the exceptional coral-like galls, and the bumpy spheres ones were restricted only to *P. palaestina* and *P. atlantica*, respectively. The surface of the galls on *P. palaestina* were smoother (no coarse galls) than those found on *P. atlantica* (Table 2A). On *P. palaestina,* 74.4% of the galls had smooth skin and 25.6% were partially crumpled. On *P. atlantica*, 42.3% were smooth, 23% were partially crumpled and 34.6% were coarse. The shape of the galls on *P. palaestina* varied, with 30.2% shaped like balls, 23.3% like flattened discuses, 27.9% like pears, and eight additional galls (18.6%), found only in one population, shaped like small “bushy corals” (see below). The shape of the galls on *P. atlantica* also varied, with 50% shaped like balls, 11.5% like flattened discuses, 34.6% like pears and 3.8% like “bumpy” spheres. Hence, neither the shape of the galls nor the surface were strictly associated with a *Pistacia* host plant species.

### 3.2 COI sequencing

Cluster analyses (Maximum Likelihood, as well as Neighbor Joining) of 630bp of the mitochondrial COI gene clearly discriminated between the *Geoica* samples from the two *Pistacia* species (Fig. 2; at least 41 SNPs; >6.5%).

**Figure 2:**
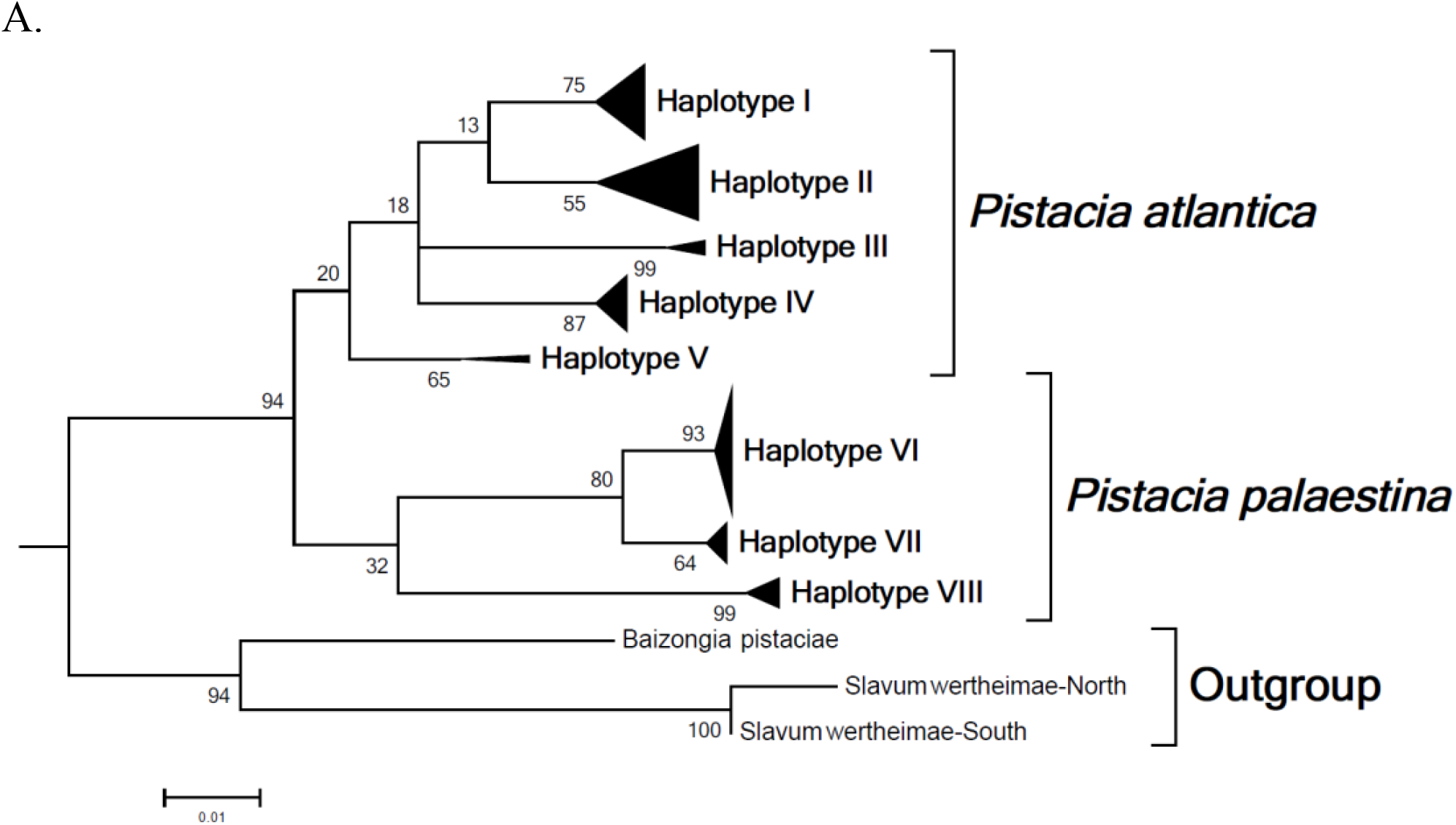
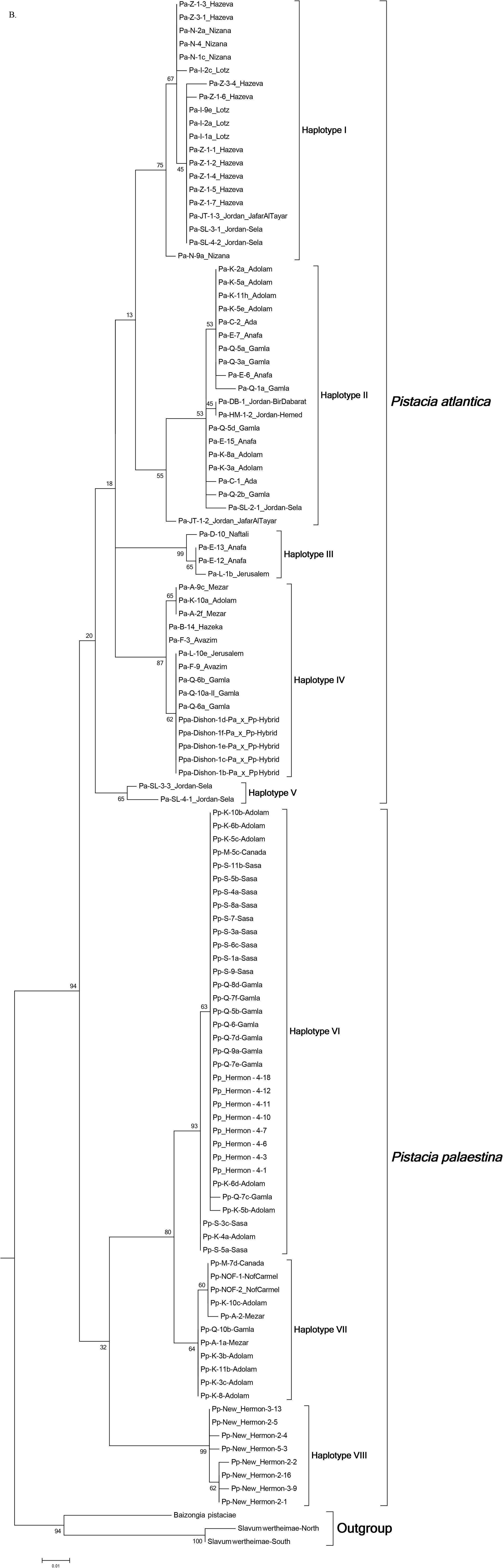
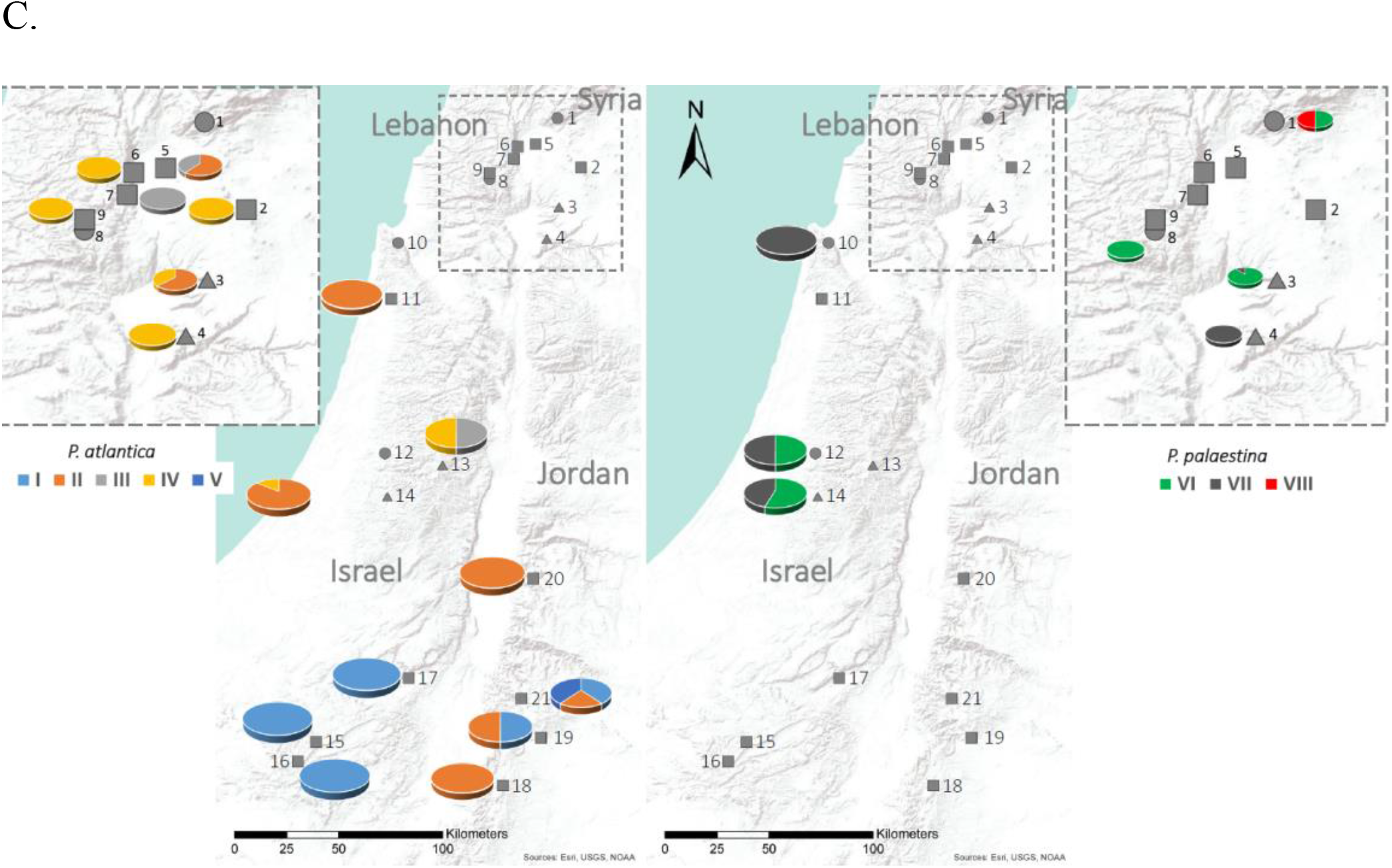
Maximum Likelihood cluster analysis, based on COI sequences, of *Geoica* spp. sampled from *P. atlantica* and *P. palaestina* in Israel and Jordan. A. A dendrogram summarizing the clustering of eight haplotypes. B. A detailed dendrogram of the various haplotypes. C. A distribution map of the various haplotypes in Israel and Jordan. Left - *P. atlantica*; Right - *P. palaestina*.

On *P. atlantica*, five *Geoica* clusters were noticeable: one haplotype (I) was found only in the xeric southern region of Israel and Jordan, and three haplotypes (II–IV) were found in the mesic central and northern regions of Israel. The fifth haplotype (V) was found only in two galls collected from Sela (Jordan). The geographic distribution of the southern haplotype (I) in Israel did not overlap with the others.

The distinction between the southern haplotype and the other three haplotypes was >26 SNPs (>4.1%). Among the three central-northern clusters, there was no indication of gall morphology, geographical or ecological differentiation, and the aphids of the various haplotypes (II–IV) were found in sympatry. The distinction between the three haplotypes was of at least 25 SNPs (>3.9%), with no indication of intermediate sequential substitutions. The Jordanian populations of Sela and Ja’far al-Tayyar harbored individuals from both the southern and the central-northern haplotypes. All five of the galls collected from the hybrid tree (*P. atlantica* x *P. palaestina*) presented haplotype IV of *Geoica* sp. of *P. atlantica*. No association was found between the various haplotypes and the morphology of the galls (Table 2B).

On *P. palaestina*, three major *Geoica* haplotypes were found (Fig. 2A). Haplotypes VI and VII differed by at least 16 SNPs (>2.5%). Both haplotypes were found sympatrically (also on the same tree) in most sampled populations, with the exception of the two northern populations, Mt. Hermon and Sasa, whose data pointed to only one haplotype of COI sequence (haplotype VI; Fig. 2B, C). No morphological association was found between the type of surface or shape of the galls and haplotypes VI or VII (Table 2B). A third haplotype (VIII) presented a distinct coral-like gall morph, differed from the other two haplotypes by at least 41 SNPs (>6.5%); it was found only in the remote area of Mt. Hermon.

### 3.3 12S sequencing

The homologies between 78 sequenced samples were generally much higher than those found for COI, and the genetic distances between samples were lower in order of magnitude (overall mean genetic distance estimation was 0.015+0.006). Nonetheless, comparable to COI division (with the same individuals expressing the equivalent haplotypes; Fig. S1), Maximum Likelihood cluster analysis of 549bp of the mitochondrial 12S gene indicated the same major four branches (haplotypes I–IV) of *Geoica* on *P. atlantica*, the two clusters on *P. palaestina* (haplotypes VI– VII) and the distinct lineage of the coral-like morph (haplotype VIII) on Mt. Hermon. The sequence differences between the *Geoica* groups on *P. atlantica* were four SNPs (0.7%), and the two subgroups of *P. palaestina* differed by only one SNP.

### 3.4 Microsatellites Analysis

A preliminary run of all tested populations and host trees included clustering possibilities for 24 potential assemblies (K values from 1 to 24; the different groups x various localities; STRUCTURE 2.3.4 admixture model; burn-in of 100,000 steps and 100,000 iterations; five replicates for each K). The results suggested differentiation between *Geoica* on *P. atlantica* and *P. palaestina* (Fig. 3A). These results were in accordance with the sequencing results of the mitochondrial COI and 12S genes. Therefore, we further considered each of these two groups on different *Pistacia* species separately.

**Figure 3:**
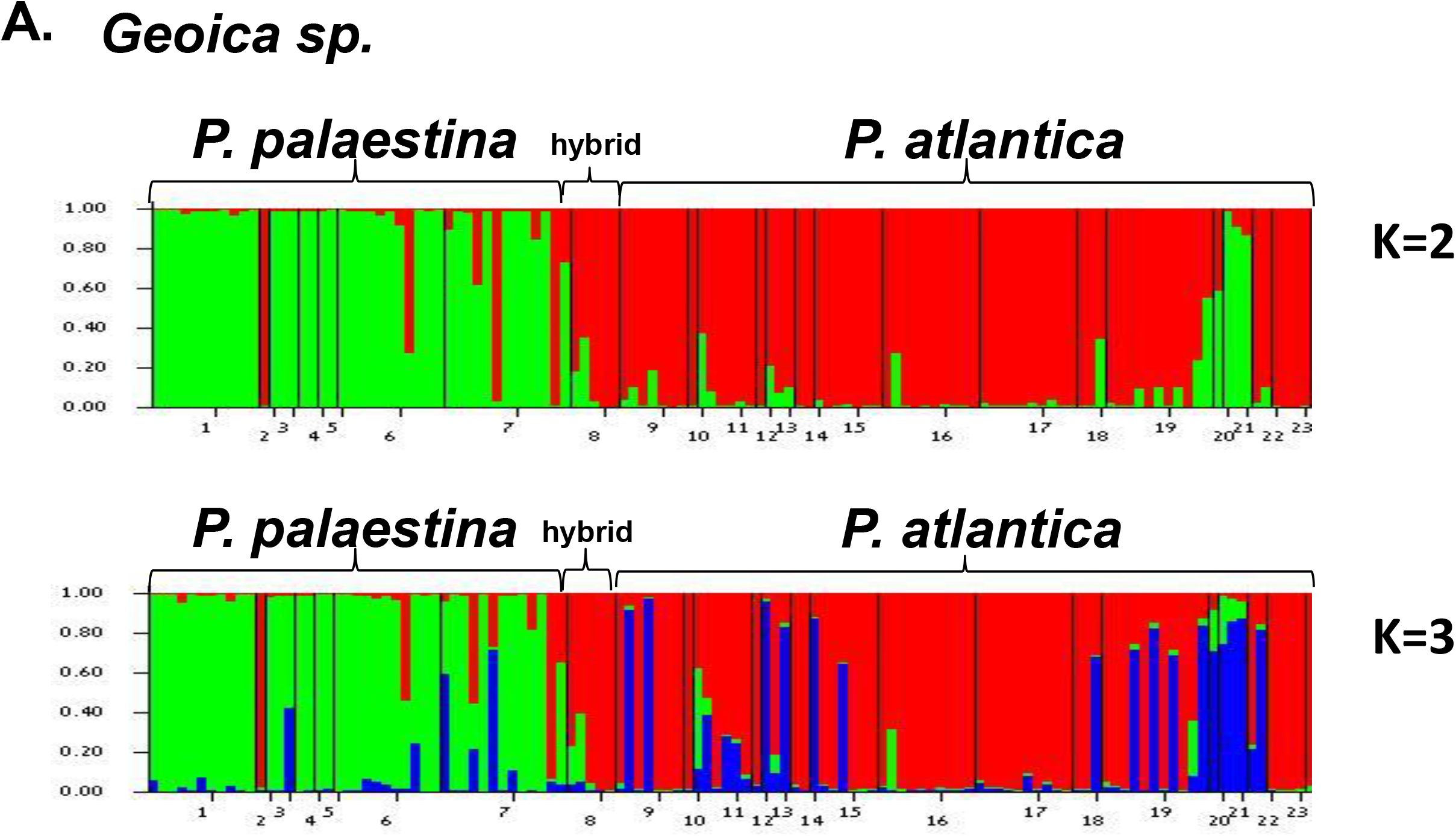
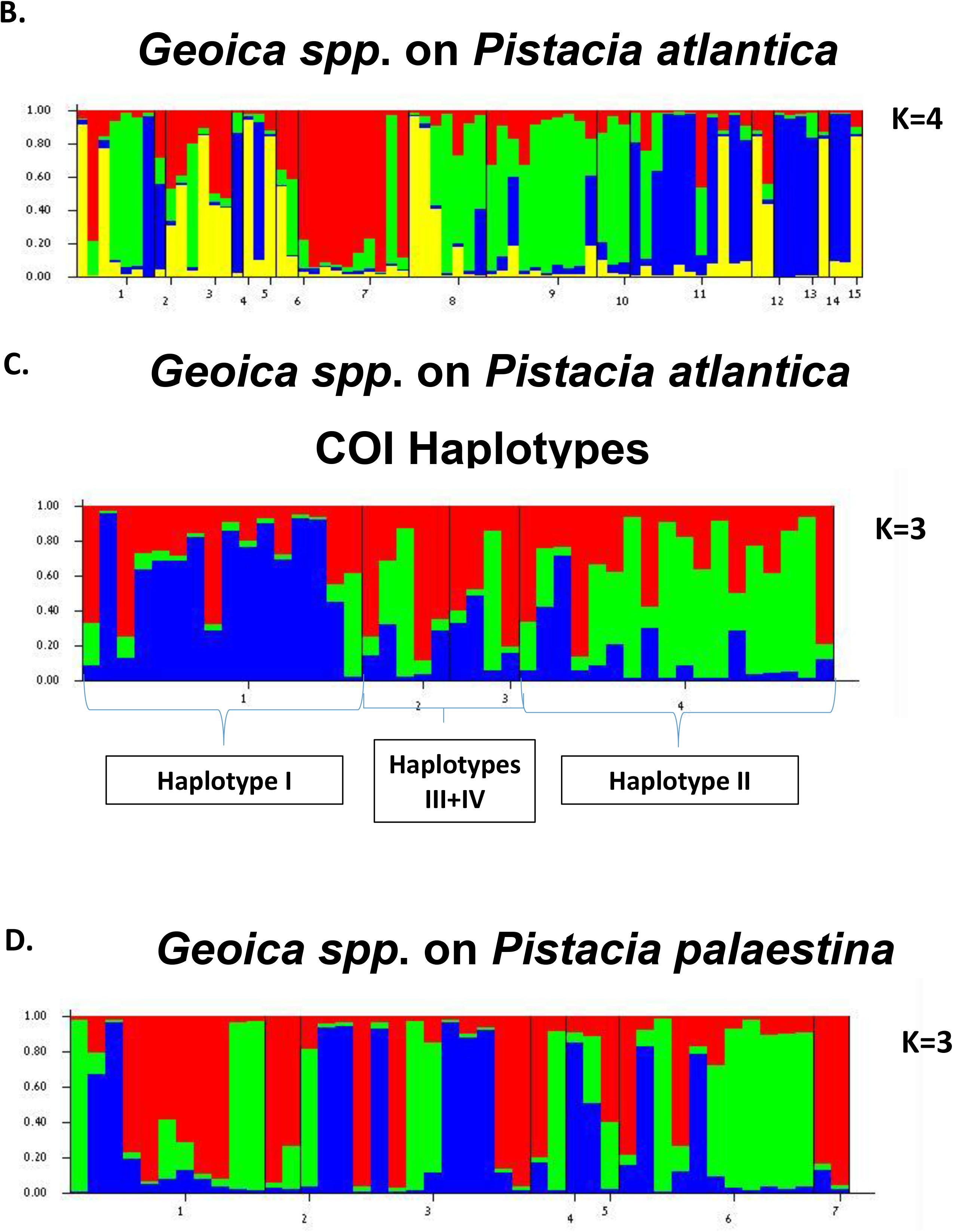
STRUCTURE Bayesian cluster analysis of microsatellites profiled under the admixture model (burn-in of 100,000 steps and 100,000 iterations; five replicates for each K). A. Differentiation between *Geoica* on *P. atlantica* and *P. palaestina (P. palaestina* — green; *P. atlantica* — red and blue; hybrid tree *P. palaestina x P. atlantica* — red as of *P. atlantica*). Populations (Pp — *P. palaestina*; Pa — *P. atlantica*). 1. Pp-Adolam; 2. Pp-Jerusalem; 3. Pp-Park Canada; 4. Pp-Nof Carmel; 5. Pp-Meizar; 6. Pp-Gamla; 7. Pp-Sasa; 8. Pp x Pa Hybrid-Dishon; 9. Pa-Gamla; 10. Pa-Meizar; 11. Pa-Tel Anafa; 12. Pa-Ein Avazim; 13. Pa-Naftali; 14. Pa-Ada; 15. Pa-Adolam; 16. Pa-Jerusalem; 17. Pa-Nizana; 18. Pa-Lotz; 19. Pa-Hazeva; 20. Pa-Bir Dabarat; 21. Pa-Ja’far al-Tayyar; 22. Pa-Hemed; 23. Pa-Sela. B. STRUCTURE differentiation (K values from 1 to 15) of *Geoica* populations on *P. atlantica*. Structure Harvester analysis indicated four different group (K=4). Populations: 1. Gamla; 2. Meizar; 3. Tel Anafa; 4. Ein Avazim; 5. Naftali; 6. Ada; 7. Adolam; 8. Jerusalem; 9. Nizana; 10. Lotz; 11. Hazeva; 12. Hemed; 13. Sela; 14. Bir Dabarat; 15. Ja’far al-Tayyar. C. STRUCTURE differentiation of the *Geoica* microsatellite profile on *P. atlantica*, according to their COI haplotypes. D. STRUCTURE differentiation (K values from 1 to 7) of *Geoica* populations on *P. palaestina.* Structure Harvester analysis indicated three different groups (K=3). Populations: 1. Gamla; 2. Meizar; 3. Sasa; 4. Nof Carmel; 5. Park Canada; 6. Adolam; 7. Jerusalem.

#### 3.4.1 Geoica sp. from P. atlantica

Mean genetic diversity parameters indicated observed heterozygosity (Ho) of 0.272+0.035 and expected heterozygosity of 0.339+0.036. The inbreeding coefficient (Fis) was positive (0.203+0.198; Table S1), but did not significantly deviate from the Hardy-Weinberg equilibrium. AMOVA analysis indicated that 35% of the variance was within individuals, 53% between individuals and 12% between populations (Table S1). The average number of migrants between populations (Nm) per generation was 1.847.

STRUCTURE and Structure Harvester analyses (100,000 Burn-in period; and 100,000 iterations; five replicates for each K; K=1-16) of 74 samples indicated clustering to four *Geoica* major subgroups (Fig. 3B). While the northern populations (1–6) showed mixed genetic types, unique profiles characterized population 7 (Adolam), the Negev Mountains populations (9 and 10), and the Negev population of Hazeva (11). The Jordanian populations (12–15) mostly showed either the genetic structure of the northern populations, or the profile of Hazeva (11). The population of Jerusalem (8) had a mixed microsatellite genetic profile resembling either the northern population or the Negev populations.

Following the COI clustering results, we further analyzed the microsatellite differentiation between the four major COI haplotypes. AMOVA showed that 10% of the molecular variance was between haplotypes (Φ_PT_=0.103; *P*=0.02). A Structure Harvester analysis (Earl and vonHoldt, 2012) indicated three major groups: one belonging to haplotype I, one to haplotype II and an additional one to haplotypes III and IV (Fig. 3C).

#### 3.4.2 Geoica sp. from P. palaestina

The mean genetic diversity parameters of samples collected throughout the *P. palaestina* distribution range in Israel revealed relatively high levels of observed and expected heterozygosity (Ho=0.461+0.054; He=0.562+0.053), with no significant deviation from the Hardy-Weinberg equilibrium (Table S2). AMOVA indicated that 42% of the variance was within individuals, 47% between individuals and 11% between populations. The average number of migrants between populations (Nm) per generation was 2.126.

STRUCTURE and Structure Harvester analyses (100,000 Burn-in period; 100,000 iterations; five replicates for each K; K=1-7) suggested three major groups (Fig. 3D). However, the STRUCTURE profile within each population was mixed. The microsatellite differentiation did not indicate any association with the two COI haplotypes.

### 3.5 The molecular clock

We tested for molecular clocks throughout the COI tree of the whole data set using the Maximum Likelihood method (Tamura and Nei, 1993; conducted in MEGA5.1), with *B. pistaciae* and *S. wertheimae*, sampled from the same trees as the *Geoica* samples, as outgroups. The analysis indicated the possibility of an equal evolutionary rate (the null hypothesis of an equal evolutionary rate throughout the tree was not rejected; *P*=1). We compared the two sequences that sowed the lowest genetic distance, one from *Geoica* found on *P. atlantica* and another on *P. palaestina* (Pa-Sela-4.1, haplotype V vs. Pp-Adolam-10c, haplotype VII), and between both sequences and the sequence of *B. pistaciae*. We found 57–58 SNPs between *Geoica* sp. and *B. pistaciae* (which parted ~34 Ma; Ren et al., 2013), and 41 SNPs between the two *Geoica* lineages. On this basis, we estimated the divergence of *Geoica* ancestries between the two *Pistacia* species as ~24-25 Ma (the Oligocene/Miocene boundary). Likewise, the divergence between the distinct coral-like gall haplotype (VIII) and the other two haplotypes on *P. palaestina* (41 SNPs) was also estimated as ~24-25 Ma. The divergence of the various haplotypes on *P. atlantica* (25–26 SNPs) was estimated as ~15 Ma (mid-Miocene), and between the two haplotypes of *P. palaestina* (16 SNPs), ~9.5 Ma.

## 4. DISCUSSION

### 4.1 Gall morphology

Gall morphological traits, although of plant tissues, are induced and largely associated with the identity of the inducing insects. The generally globular and sealed morphological appearance on the leaflet midveins of various species of *Pistacia* trees is a clear characteristic of the galls induced by *Geoica* aphids. Blackman and Eastop (2018) implied that the color, type of surface and shape of the gall slightly differ between *Pistacia* host species. We noted that the gall’s color as well as the type of skin surface are less indicative, as they may have been altered among habitats, by the gall developmental stage, and with sun exposure (Inbar and Ben-Shlomo, personal observation). As shown here (Table 2), the shape of the gall was also not a distinctive trait distinguishing between and within *P. atlantica* and *P. palaestina Geoica* galls. The delicate morphological traits (i.e., surface and spherical characteristics of the galls) most likely depend upon and are controlled by the individual *Pistacia* tree genotype or physiological machinery. Interestingly, out of eight discrete haplotypes, only one *Geoica* haplotype (*P. palaestina* — haplotype VIII) can be clearly recognized according to its distinct coral-shaped morphology (Table 2). Indeed, these narrowly distributed galling aphids in northern Israel have distinct morphological and molecular (see below) traits, which recently qualified it as a new species named *Geoica inbari* (Barjadze et al., 2022).

### 4.2 Speciation of the genus *Geoica* in the Levant

#### 4.2.1 The specificity of host-aphid interaction

In recent decades “DNA taxonomy” has become widespread in delimiting cryptic species (Fiser et al., 2018; Despres, 2019). Mitochondrial DNA, particularly a fragment of the *Cytochrome C Oxidase I* (COI) gene, is widely used as a “barcode marker” for identifying divergent lineages of insects. Attempts had been made to adopt an “objective” approach and to define species according to a sequence-divergent threshold (reviewed in Hebert et al., 2010; Krishnamurthy and Francis, 2012). The suggested rate of COI sequence-divergent threshold for insects is 3% divergent (Stoeckle and Hebert, 2008; Krishnamurthy and Francis, 2012), although there are reports of lower thresholds (e.g., several identified Lepidoptera species diverged as little as 1.3%; Hebert et al., 2010). Hence, to better assess the status of potentially cryptic species, multiple independent molecular markers are desirable (Fiser et al., 2018; Despres, 2019). Therefore, in addition to an analysis of the barcode COI marker, we analyzed a conservative mtDNA locus (12S), as well as seven known nuclear microsatellites.

The *Geoica-Pistacia* host-associated lineages clustered to entirely different clades in each and every genetic marker analyzed, both mitochondrial and nuclear. Even when both *Pistacia* species grow in sympatry, each consists solely of its own *Geoica* haplotypes. The explicit discrimination between *P. atlantica* and *P. palaestina* in both allopatric and sympatric populations suggest that the splitting of the genus *Geoica* between the two *Pistacia* host species was a single and early event. Despite the frequent landings of the sexupare on the “wrong” *Pistacia* (Wool et al., 1997), our results indicate that successful host shifts that require gall formation within this aphid complex is extremely rare, if at all.

Considering numerous molecular markers, Ren et al. (2013, 2017, 2019) estimated the divergent time of the Melaphidina, a related group of aphids (Fordini). Focusing on the gall-forming aphids on the genera *Rhus* and *Pistacia*, they suggested that the mean age of the common ancestor of the Fordinae subtribe is 74–77 Ma. They estimated that within the *Pistacia* lineage, the divergence of the Baizongiini tribe occurred no later than ~65 Ma, and the split between the genera *Baizongia* and *Geoica* took place ~34 Ma (Eocene/Oligocene boundary). Using the Ren et al. (2013) chronogram as a base time, and assuming an equal evolutionary rate, we estimated the divergent time of *Geoica* sp. lineages between the two *Pistacia* host species as 24–25 Ma (the Oligocene/Miocene boundary).

#### 4.2.2 The co-evolution Pistacia-Geoica

The origin and evolution of the genus *Pistacia* are controversial. Zohary (1952) suggested that the genus evolved ~80 Ma; Kozhoridze et al. (2015) proposed a somewhat longer evolution of ~84 Ma based on global occurrence, geostatistical analyses, and environment-based probability distribution; while Xie et al. (2014), who sequenced two nuclear and seven plastid loci of the *Pistacia* species, estimated the divergence of *Pistacia* from its closest relative at 37–38 Ma, and the divergence between *P. atlantica* and *P. palaestina* at 9.42 Ma. Our finding indicating the divergence of *Geoica* ancestries between the two *Pistacia* species at 24–25 Ma supports the longer timescale for the evolution of *Pistacia*, allowing for diversification and/or co-diversification of the host-specific *Geoica* aphids and their *Pistacia* host ancestor(s). Following the divergence between *P. atlantica* and *P. palaestina*, a further monophyletic speciation of the genus *Geoica* occurred within each *Pistacia* host lineage. The succeeding speciation of *Geoica* on *P. atlantica* was more extensive (see below); hence, considering its complex biogeographical distribution, we cannot yet determine on which *Pistacia* species the speciation of the *Geoica* genus originated.

#### 4.2.3 Genetic differentiation of Geoica spp. among and within P. atlantica populations

*P. atlantica* is distributed along the Irano-Turanian region, one of the largest floristic regions of the world. However, its evolutionary history and biogeography is less understood (reviewed in Manafzadeh et al., 2017). The collision and the deformation of Central Asian geology, as well as the complex and heterogeneous geological history, most likely vastly affect the speciation of various species inhabiting this region (Manafzadeh et al., 2017). Our results of five discrete COI haplotypes (differing by at least 25 SNPs; >3.9%), supported by the divergence of 12S, as well as the microsatellite profiles, suggest that at least five distinct *Geoica* species radiated on *P. atlantica* in the Levant. The high level of sequence divergence and the corresponding branching out of both mitochondrial and nuclear loci suggest a long-standing splitting to different varieties, estimated to take place during the mid-Miocene (~15 Ma) — a phase (Kurschner et al., 2008; Manafzadeh et al., 2017) of markedly lower the CO_2_ level, associated with both a temperature drop and major glaciation (Kurschner et al., 2008).

The heterogeneous geological history of the Irano-Turanian zone (Manafzadeh et al., 2017), together with long-lasting climatic fluctuations, triggered repeated changes in the distribution of *P. atlantica*, and consequently of *Geoica* aphids, with possible isolated populations left in refugia islands of suitable habitats (Yom-Tov and Tchernov, 1988; Danin, 1999). The lasting patchy distribution may well facilitate the accumulation of mutations, leading to further speciation of the *Geoica* complex on *P. atlantica*.

Nowadays the distribution of *P. atlantica* in Israel reflects a discontinuous spread from a mesic climate in northern and central regions to a xeric environment in the south. One of the five *Geoica* haplotypes, haplotype I, is restricted to xeric habitats. The Israeli Negev Desert is particularly dry with annual precipitation lower than 100 mm of rain (Amiran et al., 1970), and has a geographically isolated *P. atlantica* population, perhaps a relict. The distribution of the *P. atlantica* population in the Negev is currently shrinking due to increasing aridity over the last thousand years. Consequently, the populations of aphids (similar to the population of trees) are smaller and more isolated. The *Geoica* haplotype I was found only in this xeric ecoregion and no other haplotype was found in the Negev Desert. The extreme ecological conditions, in concert with the locally isolated population, are promoting rapid speciation in the Negev. Similarly, distinct allopatric genetic haplotypes and cryptic speciation were detected in the closely related gall-forming aphid *Slavum wertheimae* (Avrani et al., 2012) and *Forda riccobonii* (Ben-Shlomo and Inbar, unpublished data) on the same *P. atlantica* populations in the Negev Desert. Interestingly, *P. atlantica* does not show an equivalent genetic separation between these two ecogeographic regions (Avrani et al., 2012).

Three additional *Geoica* haplotypes were found sympatrically, also on the same tree, in the central and northern mesic regions of Israel (with over 600 mm mean annual precipitation). The existence of two *Geoica* species in northern-central Israel was noticed by Brown and Blackman (1994), who described *G. rungsi* and *G. harpazi*, which differ in their alatae female morphology. Based on the sequences of the COI and COII genes, Inbar et al. (2004) also suggested two *Geoica* species on *P. atlantica*. Unfortunately, we cannot match our results to the reported species and defined haplotypes of Brown and Blackman (1994). Notably, the two published COI sequences (AY227075 and AY227085; Inbar et al. 2004), which differ by 12 SNPs, are both clustered within the species that represents haplotype III, suggesting the possibility of yet more complex radiation. The last haplotype, V, is rare and was found only in two individuals from Sela, Jordan (population number 21, Fig. 1).

The evidently current sympatric allocation of at least three different species in northern and central Israel is probably a result of allopatric speciation, with secondary sympatry, that may have been pushed further by competition (Inbar 1998). Following the last glaciation cycle, when the climate warmed once more, the *P. atlantica* population expanded and became continuous again, bringing about the *Geoica*-galling aphids to present extensive “sympatric” allocation. The extensive and long (15 Ma) separation, together with the extensive genetic differentiation, suggest that these five haplotypes should be named as new species. Analyses of the genetic structure of *Geoica* spp. on a wider geographic scale in Central Asia and North Africa should shed light on the evolutionary history and origin of the various haplotypes on *P. atlantica*.

#### 4.2.4 Genetic variation between and within P. palaestina-Geoica populations

*P. palaestina* is widely distributed in the Mediterranean zones of Israel (Ben-Shlomo and Inbar, 2012) and is much less fragmented than the populations of *P. atlantica.* Furthermore, the distribution of this species is restricted to mesic habitats. This factor may explain why *P. palaestina* hosts fewer *Geoica* haplotypes than *P. atlantica.* Nonetheless, further sympatric speciation of *Geoica* sp. with three distinct COI haplotypes was found on *P. palaestina*. One unique *Geoica* species (*G. inbari*, haplotype VIII), which differs from the other two haplotypes by at least 41 SNPs (>6.5%), presents a clear altered form of the gall with a coral-like morphology, rather than a globular one. *G. inbari* (Haplotype VIII) puts forward an extensive ancient separation (~24–25 Ma). This distinct species is rare, and found only in the far and high Mt. Hermon.

The other two COI haplotypes differ by more than 2.8% (at least 20 SNPs) of the sequence. The 12S sequencing results showed differentiation of only one substitution, hence the allies in each group were in accordance with the COI results. While finding clear differentiation in the mitochondrial genome, the nuclear microsatellite profile of the two haplotypes looked alike. We suggest that the two haplotypes represent two cryptic species of *Geoica.* Comparable discordance between mitochondrial and nuclear diversity patterns between cryptic species are commonly found (Despres, 2019; Hinojosa et al., 2019), as the two genomes respond in a different manner to the demographic and spatial event resulting from the maternally transmitted monoploid mtDNA and the meiotic segregation and recombination of nuDNA.

One of these two species may be *G. wertheimae*, reported as the *Geoica* species found on *P. palaestina.* Three sequences of *Geoica wertheimae* are deposited in the Genbank: Accessions AH012643, AY227080, AY227090 - ‘M Wink 18450’ (Inbar et al., 2004); EU701670 - voucher CNC#HEM012658 (Foottit et al., 2008); and DQ499622 (Zhang and Qiao, 2007). The first two cluster within the lineage of *Geoica* on *P. palaestina* presented here (Fig. S2). As the sequence of Foottit et al. (2008) was identical to most samples of haplotype VI, we regard haplotype VI as *Geoica wertheimae* and suggest that haplotype VII should be named as yet another *Geoica* species.

## 5. CONCLUSIONS

The galling-aphid genus *Geoica* went through extensive radiation in the Levant. The splitting of the genus between *P. atlantica* and *P. palaestina* is estimated to have occurred 24–25 Ma (the Oligocene/Miocene boundary). Five different haplotypes suggesting five different species have been further speciating among *Geoica* spp., galling on *P. atlantica,* and an additional three species, on *P. palaestina*.

## ACKNOWLEDGEMENTS

We thank MOFET Institute for financial support and the students of Molecular Ecology workshop for their help.

## REFERENCES

Aguilar, J.M., Boecklen, W.J., 1992. Patterns of Herbivory in the *Quercus grisea* × *Quercus gambelii* Species Complex. Oikos. 64, 498–504.

Amiran, D.H.K., Elster, J., Gilead, M., Rosenman, N., Kadmon, N., Paran, U., 1970. Atlas of Israel Surveys of Israel Ministry of Labor. Elsevier, Jerusalem and Amsterdam.

Avrani, S., Ben-Shlomo, R., Inbar, M., 2012. Genetic structure of a galling aphid and its host *Pistacia atlantica* across an Irano-Turanian distribution: from fragmentation to speciation? Tree Genet. Genomes. 8, 811–820.

Barjadze, S., Halbert, S., Matile D., Ben-Shlomo, R., 2018. A new genus of gall-forming aphids of tribe Fordini Baker, 1920 (Hemiptera: Aphididae: Eriosomatinae) from Jordan. Ann. Soc. Entomol. Fr. 54, 511–521.

Barjadze, S., Halbert, S., Ben-Shlomo, R., 2022. A new gall-producing species of *Geoica* Hart, 1894 (Hemiptera: Aphididae: Eriosomatinae) from Israel. Zootaxa, in press.

Ben-Shlomo, R., Inbar, M., 2012. Patch size of gall-forming aphids: Deme formation revisited. Popul. Ecol. 54, 135–144.

Blackman, R.L., Eastop, V.F., 2018. Aphids on the world’s plants: an online identification and information guide. http://www.aphidsonworldsplants.info/

Brown, P.A., Blackman, R.L., 1994. Morphometric variation in the *Geoica utricularia* (Homoptera: Aphididae) species group on *Pistacia* (Anacardiaceae), with descriptions of new species and a key to emigrant alatae. Syst. Entomol. 19, 119–132.

Danin, A., 1999. Sandstone outcrops — A major refugium of Mediterranean flora in the xeric part of Jordan. Isr. J. Plant Sci. 47, 179–187.

Despres, L., 2019. One, two or more species? Mitonuclear discordance and species delimitation. Mol. Ecol. 28, 3845–3847.

Earl, D.A., vonHoldt, B.M., 2012. STRUCTURE HARVESTER: a website and program for visualizing STRUCTURE output and implementing the Evanno method. Conservation Genet. Resour. 4, 359–361.

Evanno, G., Regnaut, S., Goudet, J., 2005. Detecting the number of clusters of individuals using the software STRUCTURE: a simulation study. Mol. Ecol. 14, 2611–2620.

Fiser, C., Robinson, C.T., Malard, F., 2018. Cryptic species as a window into the paradigm shift of the species concept. Mol. Ecol. 27, 613–635. https://doi.org/10.1111/mec.14486.

Floate, K.D., Fernandes, G.W., Nilsson, J.A., 1996. Distinguishing intrapopulational categories of plants by their insect faunas: galls on rabbitbrush. Oecologia. 105, 221–229. https://doi.org/10.1007/BF00328550.

Foottit, R.G., Maw, H.E.L., Von dohlen, C.D., Hebert, P.D.N., 2008. DNA BARCODING — Species identification of aphids (Insecta: Hemiptera: Aphididae) through DNA barcodes. Mol. Ecol. Resour. 8, 1189–1201.

Giron, D., Huguet, E., Stone, G.N., Body, M., 2016. Insect-induced effects on plants and possible effectors used by galling and leaf-mining insects to manipulate their host-plant. J. Insect Physiol. 84, 70–89.

Golan-Goldhirsh, A., 2009. Bridging the gap between ethnobotany and biotechnology of *Pistacia*. Isr. J. Plant Sci. 57, 65–78.

Hebert, P.D.N., deWaard, J.R., Landry, J.-F., 2010. DNA barcodes for 1/1000 of the animal kingdom. Biol. Lett. 6. https://doi.org/10.1098/rsbl.2009.0848.

Hinojosa, J.C., Koubínová, D., Szenteczki, M.A., Pitteloud, C., Dincă, V., Alvarez, N., Vila, R., 2019. A mirage of cryptic species: Genomics uncover striking mitonuclear discordance in the butterfly *Thymelicus sylvestris*. Mol. Ecol. 28, 3857–3868. https://doi.org/10.1111/mec.15153.

Inbar, M., 1998. Competition, territoriality and maternal defense in a gall-forming aphid. Ethol. Ecol. Evol. 10, 159–170.

Inbar, M., 2008. Systematics of *Pistacia:* Insights from specialist parasitic aphids. Taxon. 57, 238–242.

Inbar, M., Wink, M., Wool, D., 2004. The evolution of host plant manipulation by insects: molecular and ecological evidence from gall-forming aphids on *Pistacia*. Mol. Phylogenet. Evol. 32, 504–511.

Ivens, A.B.F., Kronauer, D.J.C., Boomsma, J.J., 2010. Characterisation and cross-amplification of polymorphic microsatellite loci in ant-associated root-aphids. Conservation Genet. Resour. 3, 73–77. https://doi.org/10.1007/s12686-010-9293-3

Kafkas, S., Perl-Treves, R., 2002. Interspecific Relationships in *Pistacia* Based on RAPD Fingerprinting. HortScience. 37, 168–171. https://doi.org/10.21273/HORTSCI.37.1.168.

Koach, J., Wool, D., 1977. Geographic distribution and host specificity of gall-forming aphids (Homoptera, Fordinae) on *Pistacia* trees in Israel. Marcellia. 40, 207–216.

Kozhoridze, G., Orlovsky, N., Orlovsky, L., Blumberg, D.G., Golan-Goldhirsh, A., 2015. Geographic distribution and migration pathways of *Pistacia —* present, past and future. Ecography. 38, 1141–1154. https://doi.org/10.1111/ecog.01496.

Krishnamurthy, P. K., Francis, R.A., 2012. A critical review on the utility of DNA barcoding in biodiversity conservation. Biodivers. Conserv. 21, 1901–1919. https://doi.org/10.1007/s10531-012-0306-2.

Kurschner, W.M., Kvacek, Z., Dilcher, D.L., 2008. The impact of Miocene atmospheric carbon dioxide fluctuations on climate and the evolution of terrestrial ecosystems. Proc. Natl. Acad. Sci. U.S.A. 105, 449–453. https://doi.10.1073/pnas.0708588105.

Lymberakis, P., Poulakakis, N., Manthalou, G., Tsigenopoulos, C.S., Magoulas, A., Mylonas, M., 2007. Mitochondrial phylogeography of *Rana* (Pelophylax) populations in the Eastern Mediterranean region. Mol. Phylogenet. Evol. 44, 115–125. https://doi.org/10.1016/j.ympev.2007.03.009.

Manafzadeh, S., Staedler, Y.M., Conti, E., 2017. Visions of the past and dreams of the future in the Orient: the Irano-Turanian region from classical botany to evolutionary studies. Biol. Rev. 92, 1365–1388. https://doi.org/10.1111/brv.12287.

Michalakis, Y., Excoffier, L., 1996. A generic estimation of population subdivision using distance between alleles with special reference for microsatellite loci. Genetics. 142, 1061–1064.

Nei, M., 1978. Estimation of Average Heterozygosity and genetic distance from a small number of individuals. Genetics. 89, 583–590.

Peakall, R., Smouse, P.E., 2006. GENALEX 6: Genetic analysis in Excel. Population genetic software for teaching and research. Mol. Ecol. Notes. 6, 288–295.

Pritchard, J.K., Stephens, M., Donnelly, P., 2000. Inference of population structure using multilocus genotype data. Genetics. 155, 945–959.

Pritchard, J.K., Wen, X., Falush, D., 2010. Documentation for structure software: Version 2.3. http://pritch.bsd.uchicago.edu/structure.html.

Remaudière, G., Inbar, M., Menier, J.J., Shmida, A., 2004. Un nouveau *Geoica* gallicole sur *Pistacia atlantica* en Jordanie (Hemiptera, Aphididae, Eriosomatinae, Fordini). Revue Fr. Ent. (N.S.). 26, 37–42.

Ren, Z., Harris, A.J., Dikow, R.B., Ma, E., Zhong, Y., Wen, J., 2017. Another look at the phylogenetic relationships and intercontinental biogeography of eastern Asian-North American Rhus gall aphids (Hemiptera: Aphididae: Eriosomatinae): evidence from mitogenome sequences via genome skimming. Mol. Phylogenet. Evol. 117, 102–110.

Ren, Z., von Dohlen, C.D., Harris, A.J., Dikow, R.B., Su, X., Wen, J., 2019. Congruent phylogenetic relationships of Melaphidina aphids (Aphididae: Eriosomatinae: Fordini) according to nuclear and mitochondrial DNA data with taxonomic implications on generic limits. Plos One. https://doi.org/10.1371/journal.pone.0213181.

Ren, Z.M., Zhong, Y., Kurosu, U., Aoki, S., Ma, E.B., von Dohlen, C.D., Wen, J., 2013. Historical biogeography of eastern Asian-eastern North American disjunct Melaphidina aphids (Hemiptera: Aphididae: Eriosomatinae) on *Rhus* hosts (Anacardiaceae). Mol. Phylogenet. Evol. 69, 1146–1158.

Stoeckle, M.Y., Hebert, P.D.N., 2008. Barcode of life. Sci. Am.. 299, 82–89.

Tamura, K., Nei, M., 1993. Estimation of the number of nucleotide substitutions in the control region of mitochondrial DNA in humans and chimpanzees. Mol. Biol. Evol. 10, 512–526. https://doi.org/10.1093/oxfordjournals.molbev.a040023.

Tamura, K., Peterson, D., Peterson, N., Stecher, G., Nei, M., Kumar, S., 2011. MEGA5: molecular evolutionary genetics analysis using maximum likelihood, evolutionary distance, and maximum parsimony methods. Mol. Biol. Evol. 28, 2731–2739.

Wool, D., 2004. Galling aphids: Specialization, biological complexity, and variation. Ann. Rev. Entomol. 49, 175–192.

Wool D., Ben-Zvi, O,. 1998. Population ecology and clone dynamics of the galling aphid *Geoica wertheimae* (Sternorrhyncha: Pemphigidae: Fordinae). Eur. J. Entomol. 95, 509–518.

Wool, D., Burstein, M., 1991. A galling aphid with extra life cycle complexity: population ecology and evolutionary considerations. Res. Popul. Ecol. 33, 307–322.

Wool, D., Manheim, O., Inbar, M., 1997. Return flight of sexuparae of galling aphids to their primary host trees: implications for differential herbivory and gall (Aphidoidea: Pemphigidae: Fordinae) abundance. Ann. Entomol. Soc. Am. 90, 341–350.

Xie, L., Yang, Z.-Y., Wen, J., Li, D.-Z., Yi, T.-S., 2014. Biogeographic history of *Pistacia* (Anacardiaceae), emphasizing the evolution of the Madrean-Tethyan and the eastern Asian-Tethyan disjunctions. Mol. Phylogenet. Evol. 77, 136–146. https://doi.org/10.1016/j.ympev.2014.04.006.

Yom-Tov, Y., Tchernov, E., 1988. The Zoogeography of Israel. The distribution and abundance at a zoogeographical crossroad. W. Junk Publishers. Springer-Verlag, New York.

Zhang, H.C., Qiao, G.X., 2007. Molecular phylogeny of Fordini (Hemiptera: Aphididae: Pemphiginae) inferred from nuclear gene EF-1*a* and mitochondrial gene COI. Bull. Entomol. Res. 97, 379–386. https://doi:10.1017/S0007485307005020.

Zohary, M., 1972. *Pistacia* L. Flora Palestine. Israel Academy of Sciences and Humanities, Jerusalem. 2, 297–300.

Zohary, M., 1952. A monographical study of the genus *Pistacia*. Palest. J. Bot., Jer. Ser. 5, 187–228.

