## Supplemental Tables for "The dynamics of speciation in the gall-forming aphid *Geoica* spp. within and among *Pistacia* host tree species"

Supplement

**Table S1: Genetic diversity of *Geoica spp.* sampled from *P. atlantica***

**Grand Mean and SE over Loci and Pops**

| | Ho | uHe | Fis | Fst |
|---|---|---|---|---|
| **Mean** | 0.272 | 0.339 | 0.203 | 0.648 |
| **SE** | 0.035 | 0.036 | 0.198 | 0.074 |

**Summary AMOVA Table - *Geoica spp.* sampled from *P. atlantica***

| Source | df | SS | MS | Est. Var. | % |
|---|---|---|---|---|---|
| **Among Pops** | 12 | 75.538 | 6.295 | 0.281 | 12% |
| **Among Indiv** | 58 | 192.208 | 3.314 | 1.241 | 53% |
| **Within Indiv** | 71 | 59.000 | 0.831 | 0.831 | 35% |
| **Total** | 141 | 326.746 | | 2.353 | 100% |

| F-Statistics | Value | P(rand >= data) |
|---|---|---|
| **Fst** | 0.119 | 0.001 |
| **Fis** | 0.599 | 0.001 |
| **Fit** | 0.647 | 0.001 |
| **Nm** | 1.847 | |

**Table S2: Genetic diversity of *Geoica spp.* sampled from *P. palaestina***

**Grand Mean and SE over Loci and Pops**

| | Ho | uHe | Fis | Fst |
|---|---|---|---|---|
| **Mean** | 0.461 | 0.562 | 0.062 | 0.431 |
| **SE** | 0.054 | 0.053 | 0.139 | 0.067 |

**AMOVA Summary Table - *Geoica spp.* sampled from *P. palaestina***

| Source | df | SS | MS | Est. Var. | % |
|---|---|---|---|---|---|
| **Among Pops** | 6 | 40.482 | 6.747 | 0.276 | 11% |
| **Among Indiv** | 37 | 133.120 | 3.598 | 1.248 | 48% |
| **Within Indiv** | 44 | 48.500 | 1.102 | 1.102 | 42% |
| **Total** | 87 | 222.102 | | 2.626 | 100% |

| F-Statistics | Value | P(rand >= data) |
|---|---|---|
| **Fst** | 0.105 | 0.001 |
| **Fis** | 0.531 | 0.001 |
| **Fit** | 0.580 | 0.001 |
| **Nm** | 2.126 | |
