## Supplementary figures and images for "The dynamics of speciation in the gall-forming aphid *Geoica* spp. within and among *Pistacia* host tree species"

### Supplemental Figure 1

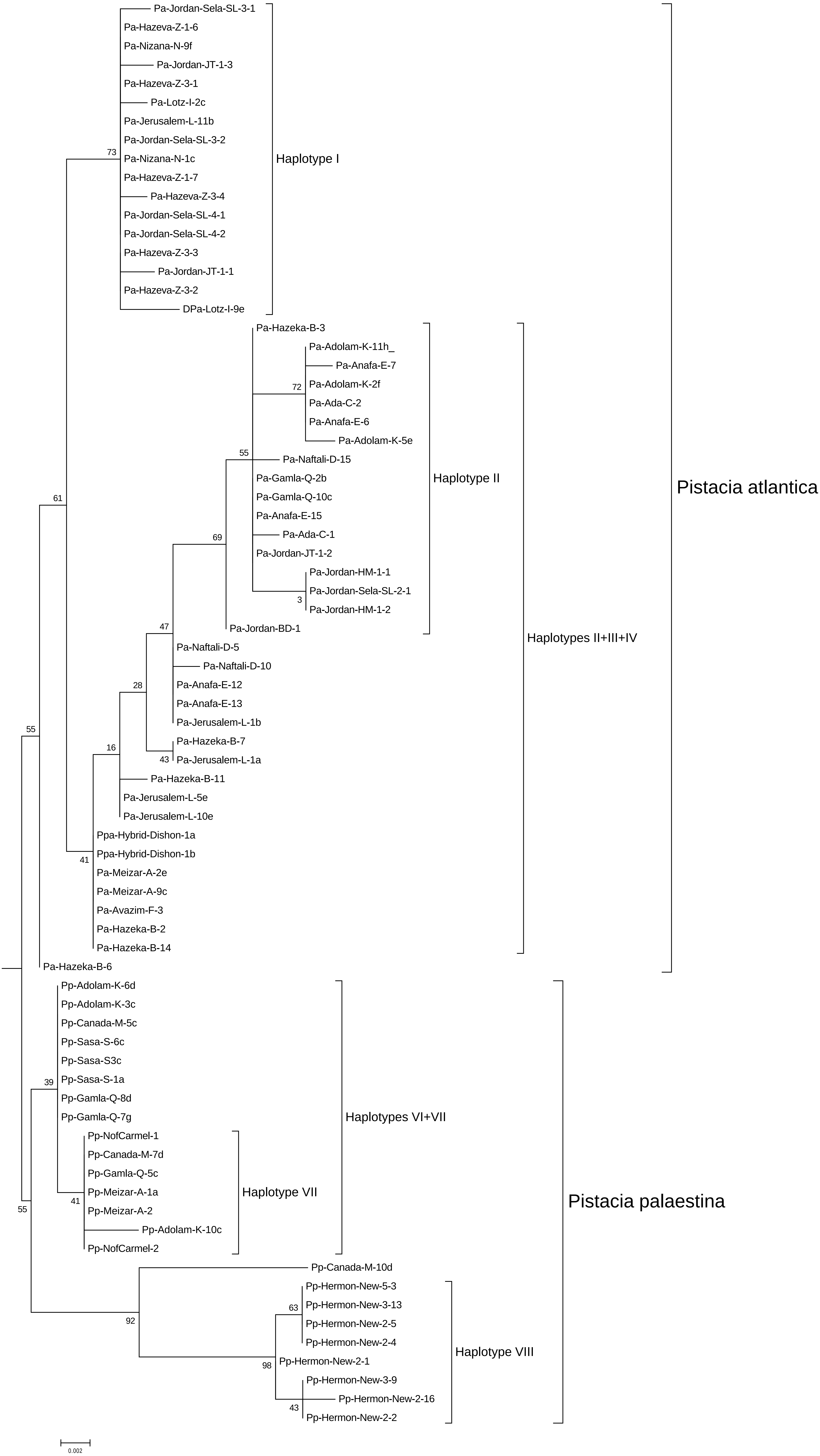

### Supplemental Figure 2

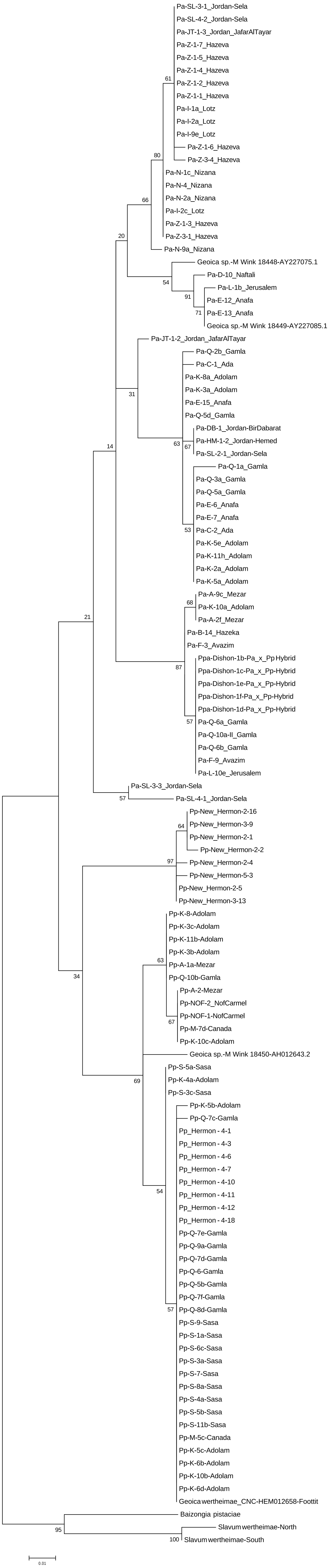
